# ppGpp Ribosome Dimerization Model for Bacterial Persister Formation and Resuscitation

**DOI:** 10.1101/663658

**Authors:** Sooyeon Song, Thomas K. Wood

**Affiliations:** Department of Chemical Engineering, Pennsylvania State University, University Park, Pennsylvania, 16802-4400, USA

**Keywords:** ppGpp, persisters, RMF, ribosomes

## Abstract

Stress is ubiquitous for bacteria and converts a subpopulation of cells into a dormant state known as persistence, in which cells are tolerant to antimicrobials. These cells revive rapidly when the stress is removed and are likely the cause of many recurring infections such as those associated with tuberculosis, cystic fibrosis, and Lyme disease. However, how persister cells are formed is not understood well. Here we propose the ppGpp ribosome dimerization persister (PRDP) model in which the alarmone guanosine pentaphosphate/tetraphosphate (henceforth ppGpp) generates persister cells directly by inactivating ribosomes via the ribosome modulation factor (RMF) and the hibernation promoting factor (Hpf). We demonstrate both RMF and HpF increase persistence and reduce single-cell persister resuscitation and that ppGpp has no effect on single-cell persister resuscitation. Hence, a direct connection between ppGpp and persistence is shown.

## INTRODUCTION

Persisters may are formed from myriad stresses such as nutrient, antibiotic, and oxidative stress (*1, 2*). In this resting state, persisters survive stresses not by actively combatting the stress but by sleeping through it (*3*). We have shown that nutrient stress creates persister cells that are equivalent to the cells known as viable but non-culturable (*2*). Since, nearly all tested cells form persisters (*4*) and almost all cells will face starvation, the persister state is probably an universal resting state (*5*). Unfortunately, since persister cells usually exist as a sub-population of less than 1% of cells, they have been difficult to study and little is known about how they form.

By inhibiting the activity of ribosomes, we have developed means to convert *E. coli* cells into a population that consists solely of persister cells (*6*) so that insights may be made about persister cell resuscitation. Using this approach, which has been verified by six independent labs to produce *bona fide* persister cells (*7–12*), we have shown that persister cells resuscitate rapidly primarily in response to external signals (*13*), such as fresh nutrients, rather than stochastically (*14*). Critically, persister cell resuscitation is heterogeneous and depends on the number of active ribosomes; once a threshold of active ribosomes is reached, persister cells elongate and divide (*13*). We also determined that persister cells sense external nutrients via their chemotaxis (for amino acids) and phosphotransferase membrane proteins (for glucose) (*14*) rather than utilizing specialized proteins for resuscitation. These external signals are propagated internally from the membrane sensors by reducing cAMP which leads to the rescue of stalled ribosomes and to the dissociation of inactive dimerized 100S ribosomes (*14*). Furthermore, resuscitating cells undergo chemotaxis toward fresh nutrients since it was the lack of nutrients that caused the persistence in the first place (*14*).

In contrast to how persister cells revive, many questions remain about how persister cells form. It is clear that inhibiting translation by stopping transcription, corrupting ribosomes, and depriving ribosomes of ATP results in a 10,000-fold increase in persistence (*6*), so inactivation of ribosome activity is imperative for persistence. This is logical in that cells are roughly 50% protein and over 90% of ATP is used for protein synthesis (*15*), so inhibiting translation effectively makes cells dormant.

Beyond translation inactivation, there is consensus for a role of ppGpp in the formation of persister cells (*16*) since ppGpp is required for high persister levels in *E. coli* (*17, 18*) and *Pseudomonas aeruginosa* (*19*). In *E. coli*, ppGpp is formed in the stationary phase in response to stress such as nutrient depletion (specifically, due to a lack of charged tRNAs); ppGpp is synthesized by RelA and SpoT (which can also degrade ppGpp) and serves to reduce ribosome synthesis (*15*) by directly modulating transcription via its interaction with RNA polymerase, via activation of RpoS (sigma^S^, the stress response sigma factor for the stationary phase), and via activation of RpoE (sigma^E^, the stress response sigma factor for misfolded proteins in the periplasm) (*20*). Critically, along with re-directing transcription, the alarmone reduces translation by activating the expression of the small (55 aa) ribosome modulation factor (RMF) (*21*), which converts active 70S ribosomes into inactive 100S ribosomes (*22*) via an inactive 90S dimer complex (*23*). This dimerization is essential for survival in the stationary phase (*24*). Also, ppGpp induces the hibernation promoting factor Hpf (95 aa) which converts 90S ribosomes into 100S ribosomes and is highly conserved as it is found in most bacteria and some plants (*25*).

Since overproduction of toxins of toxin/antitoxin (TA) systems and production of any toxin increases persistence (*18*) and since ppGpp is necessary for full activity of toxins MazF (*26*) and HipA7 (*17*), the prevailing model is that ppGpp activates TA system toxins (*27*). In a now defunct model (*28*), the Gerdes group proposed that persisters cells were formed through ppGpp based on its activation of Lon protease (via polyphosphate); Lon would then degrade preferentially antitoxins so that toxins would be activated. Here we propose a simpler model: ppGpp induces persistence directly through ribosome dimerization, and persister cell resuscitation is through reactivation of ribosomes. Hence, we provide the long-sought connection between ppGpp and persistence as well as show RMF is important not only for the stationary phase but also for persistence.

## RESULTS

We discovered that the protein that activates translation by dissociating hibernating 100S ribosomes, the GTPase HflX (*29, 30*), significantly increases the rate persister cells resuscitate and that the *hflX* mutation prevented resuscitation (*14*). Therefore, we hypothesized that dimerization of ribosomes by RMF causes persistence and reactivation of 100S ribosomes resuscitates persister cells. It is well-established that (i) ppGpp is formed during myriad stresses such as depletion of nutrients, phosphate, fatty acids, and iron as well as due to osmotic and acid stress (*20*) so ppGpp is utilized by stressed cells, (ii) ppGpp induces *rmf* (*21*), (iii) RMF dimerizes and inactivates ribosomes (*22*), and (iv) ppGpp increases persistence (*16–19*). To complete our model, we investigated the role of RMF in forming persister cells and their resuscitation. We also tested the impact of the 100S dimerization factor Hpf, that converts 90S ribosome into 100S ribosomes (*25*) and ppGpp on persister cell resuscitation.

### RMF and Hpf increase persistence

If RMF increases persistence, then deleting *rmf* should reduce the number of persister cells with antibiotics. Using ampicillin at 10 MIC (100 µg/mL), we found there was a 1,480-fold reduction in the number of persister cells (**Fig. 1A**). Similarly, inactivating Hpf resulted in a 12-fold reduction in persistence with ampicillin. Therefore, RMF is more important than Hpf for forming persister cells. In addition, as expected, overproducing both RMF and Hpf reduced single-cell resuscitation, but only moderately (∼20%) (**Fig. 1B**).

**Fig. 1.**
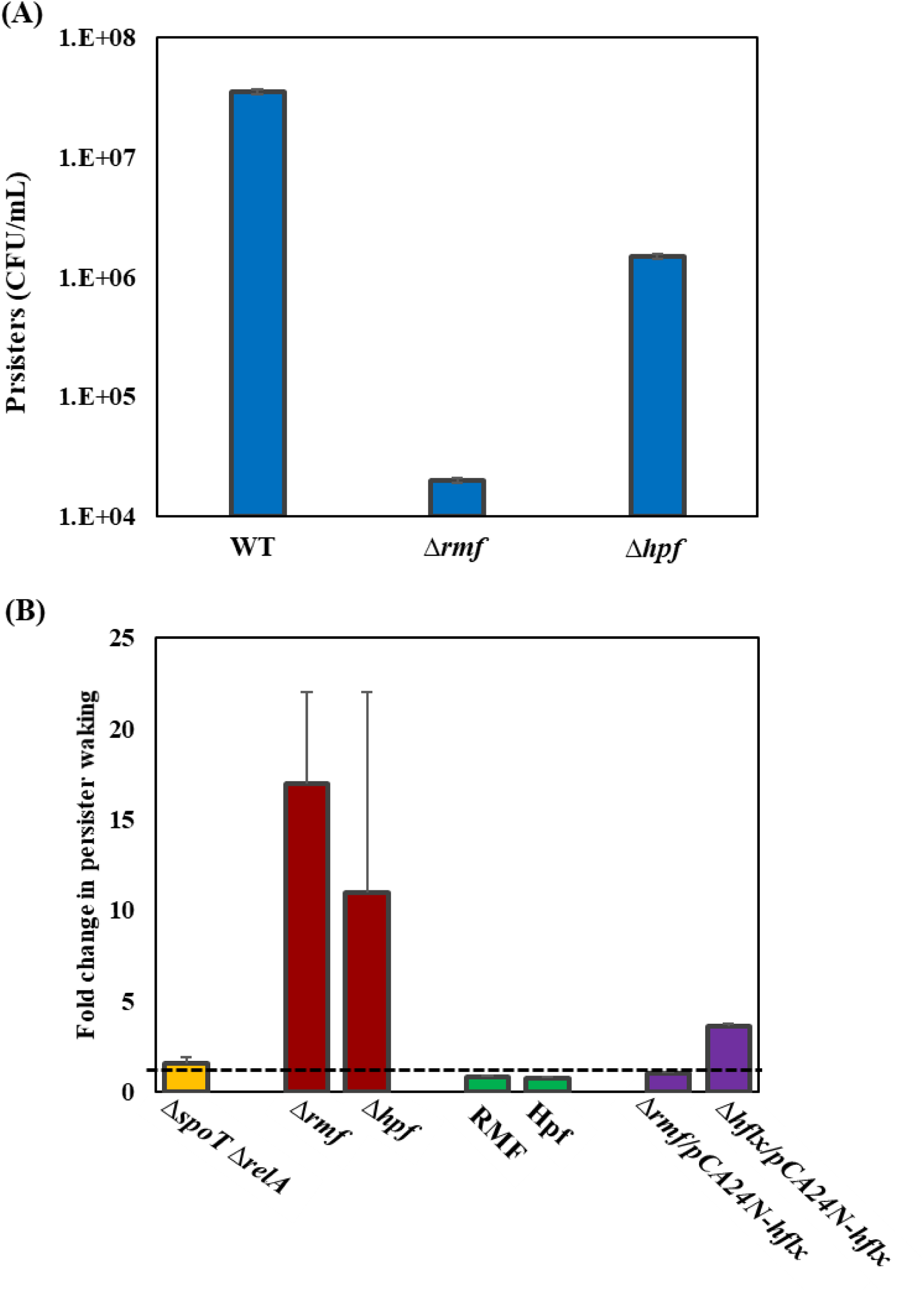
Ribosome-associated factors RMF and Hpf increase persistence and reduce resuscitation. **(A)** Persister cell formation of the isogenic mutants Δ*rmf*, and Δ*hpf* after treating with ampicillin for 3 h at 100 µg/mL. The results are from two independent experiments. **(B)** Single-cell persister resuscitation as determined using light microscopy (Zeiss Axio Scope.A1). The total and waking number of persister cells are shown in **Table S2**. Microscope images are shown in **Fig. S1, S2, S3** and **S4**. The fold-change in resuscitation is relative to MG1655 for MG1655 Δ*spoT* Δ*relA*, relative to BW25113 for the BW25113 deletion mutants, relative to BW25113/pCA24N for strains producing proteins from pCA24N plasmids in BW25113, and relative to Δ*rmf/*pCA24N*-hflx* for Δ*hflx/*pCA24N*-hflx*. M9 glucose (0.4%) agarose gel pads were used for all the strains except MG1655 and its isogenic mutant Δ*spoT* Δ*relA* since the Δ*spoT* Δ*relA* mutant cannot grown on minimal medium; hence, LB agarose gel pads were used for these two strains. The results are the combined observations from two independent experiments after 3 h for the MG1655 and Δ*spoT* Δ*relA*, after 4 h for BW25113 and its deletion mutants, and after 6 h for cells harboring pCA24N and its derivatives. For both panels, error bars indicate standard deviations.

### ppGpp doe s not affect waking

Although ppGpp is the single-most important factor for forming persisters, its role in resuscitation in not clear. To investigate this at the single cell level in a manner that allows us to capture initial waking events (*13*, 14), we converted the complete population into persister cells (*6*), a process that has been confirmed eight ways (*13*), and investigated single-cell resuscitation in the absence of ppGpp via the *relA spoT* mutant. We utilized rich medium since the *relA spoT* mutant does not grow on minimal medium and found there was little effect of eliminating ppGpp on single-cell persister resuscitation (**Fig. 1B**) although reviving *relA spoT* cells were clearly elongated (**Fig. S1)**. These results agree with those found for resuscitating starved *P. aeruginosa* on agar plates where *relA spoT* had no effect (*33*).

### RMF and Hfp reduce single-cell waking

We hypothesized that inactivating dimerization factors RMF and Hpf would lead to faster persister cell resuscitation since ribosomes could not be dimerized during persister cell formation; hence, we quantified single cell resuscitation for the isogenic *rmf* mutant. In agreement with the hypothesis, we found cell resuscitation with minimal glucose medium was increased dramatically (17-fold) (**Fig. 1B**). Corroborating our model, the isogenic *hpf* mutant also increased waking dramatically (12-fold) (**Fig. 1B**). Hence, RMF and Hpf reduce persister waking by maintaining ribosomes in their inactive state.

### HflX increases waking only in the presence of RMF

Since inactivating HflX, which dissociates hibernating 100S ribosomes to resuscitate persister cells, prevents cell resuscitation (*14*), we hypothesized that HflX would only increase waking in a background where RMF was used to induce dormancy (by forming inactive 100S ribosomes). Therefore, we compared producing HflX in the *rmf* mutant vs. producing HflX in a *hflX* mutant (so there was no background HflX from the chromosome but there was RMF to dimerize ribosomes) and found there was a four-fold increase when RMF was present (**Fig. 1B**). Therefore, HflX increases waking when ribosomes are dimerized by RMF.

## DISCUSSION

Since we have shown previously that the protein that reactivates translation by dissociating 100S hybridized ribosomes, HflX, is important for single-cell persister resuscitation (*14*), it is logical that hybridization of ribosomes by RMF should be required for the formation of persister cells. Hence, our ppGpp ribosome dimerization persister (PRDP) model for persister cell formation and resuscitation focuses on ribosome inactivation and activation, respectively (**Fig. 2**). Critically, 100S ribosome dissociation and recovery is extremely rapid (less than one minute) (*25*) which allows for rapid persister cell resuscitation; this matches our results where we found the resuscitation of single persister cells is rapid and the speed depends on the number of active ribosomes (*13*). Furthermore, since RMF is conserved in bacteria (*25*), our results suggest the PRDP model be general for the formation of the persister state. Also, since our results showing RMF increases persistence with the antibiotic ampicillin are consistent with previous results showing that RMF increases tolerance to netilmicin (*34*), gentamicin (*35*), acid (*32*), osmotic stress (*36*) and nutrient limitation in the stationary phase (*37*, 38), the PRDP model has general applicability for explaining how myriad stresses lead to persister cell formation as well as for explaining how persister cells resuscitate from these myriad resting states.

**Fig. 2.**
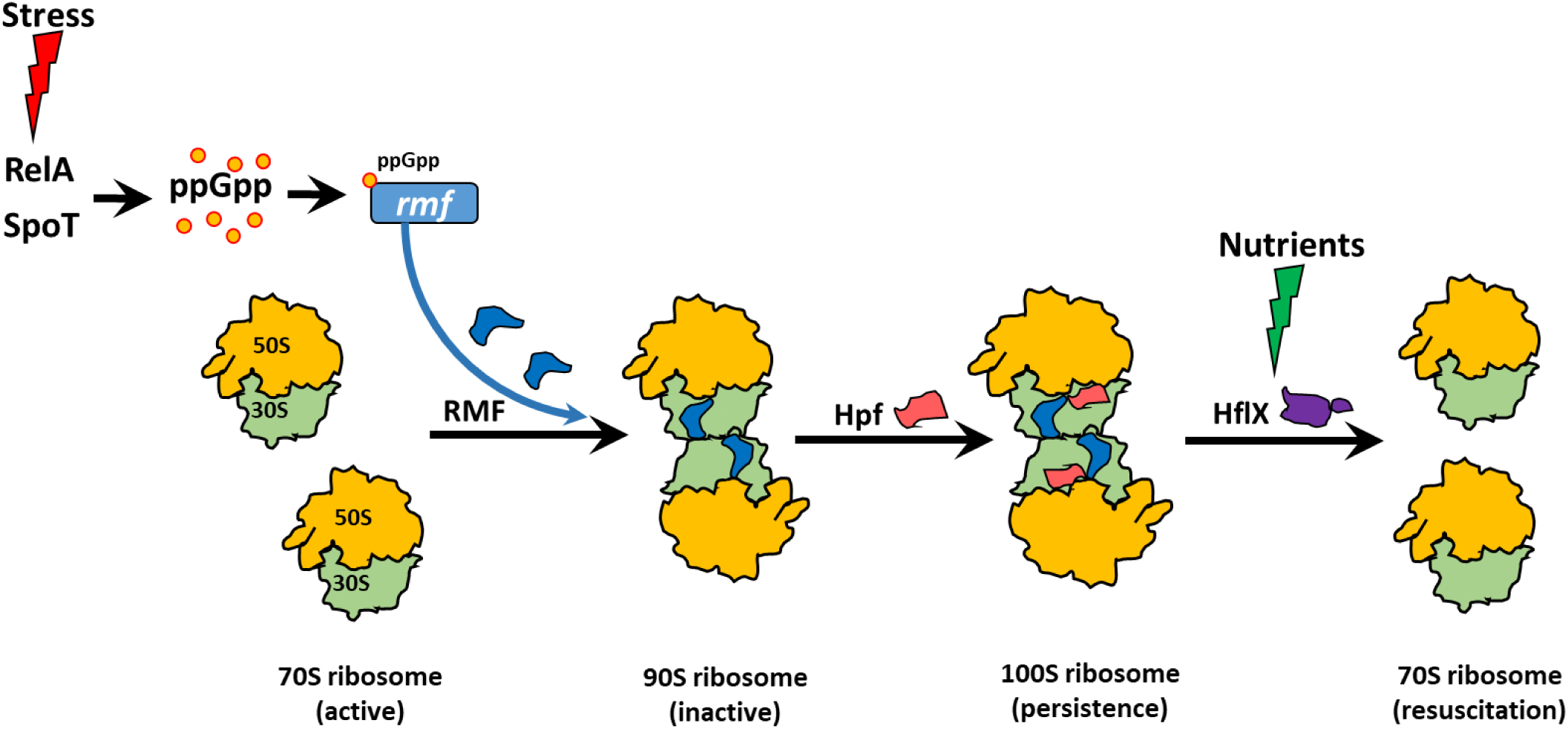
Schematic of the ppGpp ribosome dimerization persister model. Myriad stresses (e.g., nutrient limitation, osmotic, acid) induce the stringent response which results in ppGpp formation by RelA/SpoT. ppGpp induces production of both RMF and Hpf which converts active 70S ribosomes into inactive 90s and 100s ribosomes, respectively. Upon addition of nutrients and removal of the stress, HflX dissociates 100S ribosomes into active 70S ribosomes and growth resumes.

Along with RMF, we found converting 90S ribosomes into 100S ribosomes via Hpf is important for increasing persistence and inactivating Hpf increases waking. Our results differ from those of *P. aeruginosa* since, in *E. coli*, inactivating Hpf increases waking dramatically (**Fig. 1B**) whereas in *P. aeruginosa*, Hpf is important for both entering dormancy and resuscitation (*33*). However, both our work and that with *P. aeruginosa* demonstrate the link of ribosomes with persistence, but our work discovers the direct link between the stringent response (i.e., ppGpp), persistence, and resuscitation.

With this direct link between ppGpp and persistence, our PRDP model bypasses the need for ppGpp to activate toxins of TA systems for the cell to become persistent. Unfortunately, the main links between TA systems and persistence are based on use of the HipA7 variant that lacks toxicity (*17, 39*). Instead, the roles of TA systems appear now to be primarily phage inhibition (*40*), plasmid maintenance, stress response, and biofilm formation (*5*). This fits well with recent results that diminish the role for TA systems in forming persister cells (*31*) and instead emphasize the role of ppGpp (*16*). Therefore, RMF and Hpf catalyzing ribosome inactivation through the formation of 100S ribosomes is the elusive connection between the stringent response and persistence.

## MATERIALS AND METHODS

### Bacteria and growth conditions

The strains (**Table S3**) were grown routinely in lysogeny broth (*41*) at 37°C. For overnight cultures, chloramphenicol (30 μg/mL) was used to retain pCA24N-based plasmids (*42*) and kanamycin (50 μg/mL) was used for deletion mutants.

### Persister cells

Persister cells were generated (*6, 13*) from exponentially-growing cells (turbidity of 0.8 at 600 nm) by adding rifampicin (100 µg/mL) for 30 min to stop transcription, centrifuging, and adding LB with ampicillin (100 µg/mL) for 3 h to lyse any non-persister cells. Cell pellets were washed twice with 0.85% NaCl then re-suspended in 0.85% NaCl.

### Persister cell assays

To determine the number of persister cells after antibiotic treatment, exponentially-growing cells (turbidity of 0.8 at 600 nm) were treated with ampicillin (100 µg/mL) for 3 h, washed twice with 0.85% NaCl to remove the ampicillin, diluted in 0.85% NaCl, and the number of persister cells was determined via a drop assay (*43*).

### Single-cell persister resuscitation

Briefly, cell populations (5 µL) of 100% persister cells were added to 1.5% agarose gel pads containing either LB or M9 glucose (0.4 wt%) medium (*44*) as indicated, and resuscitation was monitored at 37°C via a light microscope (Zeiss Axio Scope.A1, bl_ph channel at 1000 ms exposure). At least two independent cultures were used at with two different positions per culture (ca. 150 to 300 individual cells were monitored).

## ACKNOWLEDGEMENTS

This work was supported by funds derived from the Biotechnology Endowed Professorship at the Pennsylvania State University. The authors have no conflicts of interest.

## Supporting Information

**Table S1.** RMF and Hpf increase persistence. Persister cells were assayed after treating with ampicillin (Amp) for 3 h at 100 µg/mL; 0 h indicates the initial cell population before ampicillin treatment. Fold-changes are relative to that of wild-type BW25113. The results are from two independent experiments.

|  | **0 h**<br>**(CFU/mL)** | **Amp**<br>**(CFU/mL)** | **Persisters**<br>**(%)** | **Fold-change** |
| --- | --- | --- | --- | --- |
| **BW25113 WT** | $7.8 \pm 0.7 \times 10^8$ | $35 \pm 6 \times 10^6$ | $4.5 \pm 0.4$ | - |
| *Δrmf* | $6.5 \pm 0.7 \times 10^8$ | $2.0 \pm 0.9 \times 10^4$ | $0.003 \pm 0.002$ | $-1,517 \pm 760$ |
| *Δhpf* | $4.0 \pm 0.2 \times 10^8$ | $15 \pm 3 \times 10^5$ | $0.37 \pm 0.07$ | $-12 \pm 3$ |

**Table S2.** Single persister cell resuscitation. Single-cell persister resuscitation as determined using light microscopy (Zeiss Axio Scope.A1). The fold-change in resuscitation is relative to MG1655 for MG1655 Δ*spoT* Δ*relA*, relative to BW25113 for the BW25113 deletion mutants, relative to BW25113/pCA24N for strains producing proteins from pCA24N plasmids in BW25113, and relative to Δ*rmf/*pCA24N*-hflx* for Δ*hflx/*pCA24N*-hflx*. M9 glucose (0.4%) agarose gel pads were used for all the strains except MG1655 and its isogenic mutant Δ*spoT* Δ*relA* since the Δ*spoT* Δ*relA* mutant cannot grown on minimal medium; hence, LB agarose gel pads were used for these two strains. The results are the combined observations from two independent experiments after 3 h for the MG1655 and Δ*spoT* Δ*relA*, after 4 h for BW25113 and its deletion mutants, and after 6 h for cells harboring pCA24N and its derivatives. The microscope images are shown in **Fig. S1, S2, S3**, and **S4**. Standard deviations are shown, and each strain was visualized at 14 positions.

|  | Total cells | Waking cells | % waking | Fold-change |
| --- | --- | --- | --- | --- |
| **MG1655 WT** | 95 ± 35 | 12 ± 3 | 13 ± 2 | 1 |
| **MG1655 $\Delta spoT \Delta relA$** | 59 ± 30 | 12 ± 4 | 21 ± 5 | 1.6 ± 0.3 |
| **BW25113 WT** | 116 ± 46 | 2 ± 2 | 2 ± 1 | 1 |
| *$\Delta rmf$* | 118 ± 35 | 40 ± 16 | 33 ± 3 | 17 ± 5 |
| *$\Delta hpf$* | 290 ± 20 | 68 ± 57 | 23 ± 20 | 11 ± 11 |
| **pCA24N** | 252 ± 20 | 24 ± 7 | 9.5 ± 2 | 1 |
| **pCA24N-*rmf*** | 123 ± 41 | 10 ± 2 | 7.9 ± 0.9 | -1.20 ± 0.07 |
| **pCA24N-*hpf*** | 392 ± 210 | 29 ± 15 | 7.28 ± 0.05 | -1.30 ± 0.06 |
| **$\Delta rmf$/pCA24N-*hflx*** | 162 ± 10 | 11 ± 5 | 6 ± 3 | 1 |
| **$\Delta hflx$/pCA24N-*hflx*** | 383 ± 18 | 88 ± 8 | 23 ± 1 | 3.6 ± 0.2 |

**Table S3.** Bacterial strains and plasmids used in this study.

| Strains | Features | Source |
| --- | --- | --- |
| *E. coli* BW25113 | Wild type | (45) |
| *E. coli* BW25113 $\Delta rmf$ | $\Delta rmf$, Km<sup>R</sup> | (45) |
| *E. coli* BW25113 $\Delta hpf$ | $\Delta hpf$, Km<sup>R</sup> | (45) |
| *E. coli* BW25113 $\Delta hflx$ | $\Delta hflX$, Km<sup>R</sup> | (45) |
| *E. coli* MG1655 | Wild type | (46) |
| MG1655 $\Delta spoT \Delta relA$ | $\Delta spoT \Delta relA$ | (47) |
| **Plasmids** |  |  |
| pCA24N | Cm<sup>R</sup>, *lacI<sup>q</sup>* | (42) |
| pCA24N_*rmf* | Cm<sup>R</sup>, *lacI<sup>q</sup>*, P<sub>T5-lac</sub>::*rmf*<sup>+</sup> | (42) |
| pCA24N_*hpf* | Cm<sup>R</sup>, *lacI<sup>q</sup>*, P<sub>T5-lac</sub>::*hpf*<sup>+</sup> | (42) |
| pCA24N_*hflX* | Cm<sup>R</sup>, *lacI<sup>q</sup>*, P<sub>T5-lac</sub>::*hflX*<sup>+</sup> | (42) |

Km<sup>R</sup> and Cm<sup>R</sup> indicate kanamycin and chloramphenicol resistance, respectively.

**Supplementary Figure 1.**
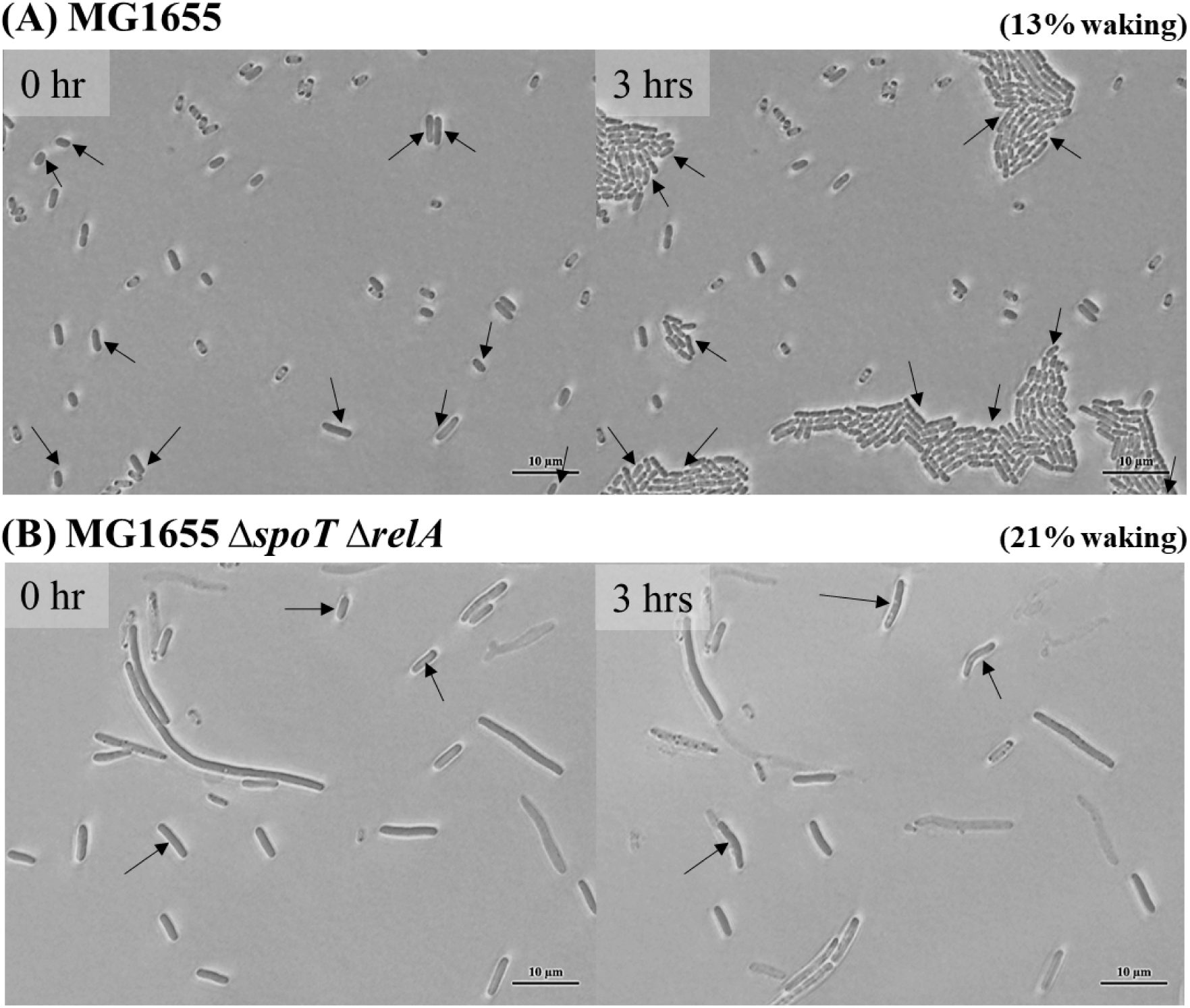
Single persister cell waking on LB after inactivating SpoT and RelA. Persister cell resuscitation for (**A**) MG1655 and (**B**) MG1655 δ*spoT* δ*relA* on LB agarose gel pads after 3 h at 37°C (note the δ*spoT* δ*relA* mutant cannot grow on minimal medium). Black arrows indicate cells that resuscitate. Scale bar indicates 10 µm. Representative results from two independent cultures are shown. Cell numbers are shown in **Table S2**.

**Supplementary Figure 2.**
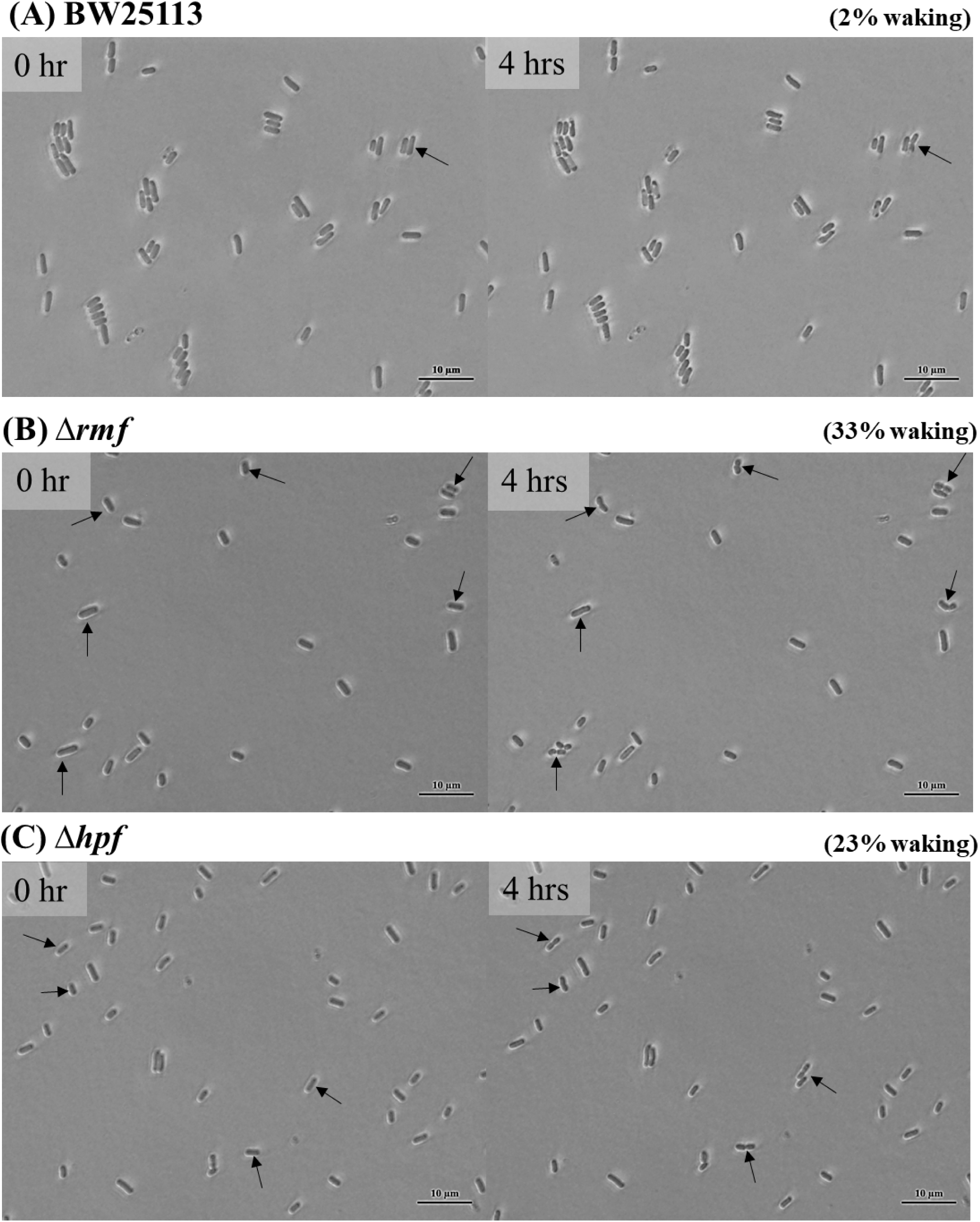
Single persister cell waking on M9 glucose (0.4%) after inactivating RMF and Hpf. Persister cells of (**A**) BW25113, (**B**) BW25113 δ*rmf*, and **(C)** BW25113 δ*hpf* waking on M9 glucose (0.4%) agarose gel pads after 4 h at 37°C. Black arrows indicate cells that resuscitate. Scale bar indicates 10 µm. Representative results from two independent cultures are shown. Cell numbers are shown in **Table S2**.

**Supplementary Figure 3.**
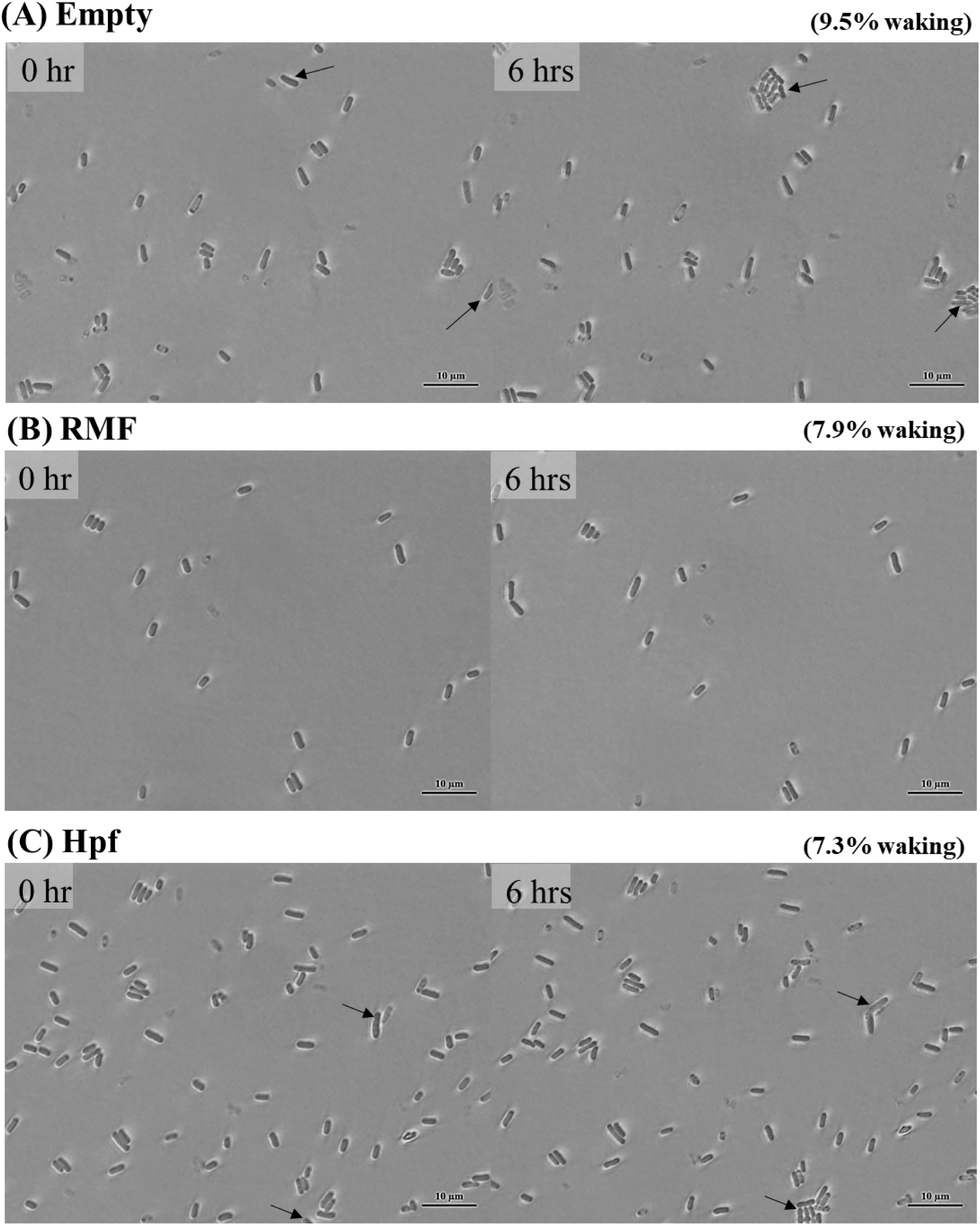
Single persister cell waking on M9 glucose (0.4%) after producing RMF and Hpf. Persister cells of (**A**) BW25113/pCA24N (“Empty”), (**B**) BW25113/pCA24N_*rmf*, and **(C)** BW25113/pCA24N_*hpf* waking on M9 glucose (0.4%) agarose gel pads after 6 h at 37°C. Black arrows indicate cells that resuscitate. Scale bar indicates 10 µm. Representative results from two independent cultures are shown. Cell numbers are shown in **Table S2**.

**Supplementary Figure 4.**
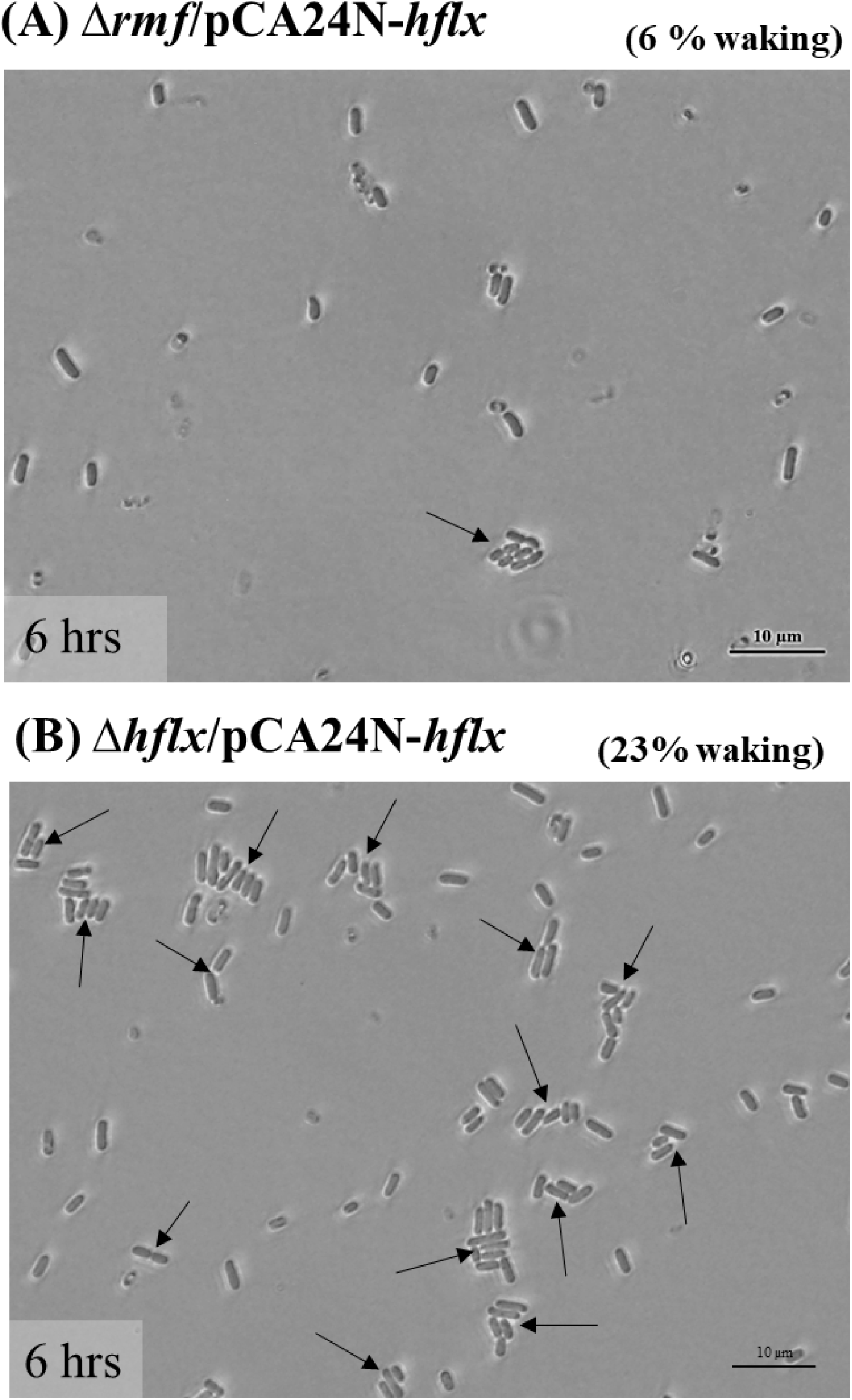
Single persister cell waking on M9 glucose (0.4%) after producing HflX. Persister cells of (**A**) Δ*rmf*/pCA24N_*hflx*, (**B**) Δ*hflx*/pCA24N_*hflx* on M9 glucose (0.4%) agarose gel pads after 6 h at 37°C. Black arrows indicate cells that resuscitate. Scale bar indicates 10 µm. Representative results from two independent cultures are shown. Cell numbers are shown in **Table S2**.

